# The Topology of Representational Geometry

**DOI:** 10.1101/2024.02.16.579506

**Authors:** Shael Brown, Reza Farivar

**Author notes:** S.B. and R.F. designed research; S.B. performed research; R.F. supervised; and S.B. and R.F. wrote the paper. The authors declare that there are no competing interests.

## Abstract

Representational similarity analysis (RSA) is a powerful tool for abstracting and then comparing neural representations across brains, regions, models and modalities. However, typical RSA analyses compares pairs of representational dissimilarities to judge similarity of two neural systems, and we argue that such methods can not capture the shape of representational spaces. By leveraging tools from computational topology, which can probe the shape of high-dimensional data, we augment RSA to be able to detect more subtle yet real differences and similarities of representational geometries. This new method could be used in conjunction with regular RSA in order to make new inferences about neural function.

**Significance Statement:** Big data in high-dimensional spaces, like neuroimaging datasets, contain important shape structures. These shape structures can be analyzed to identify the underlying features and dynamics which drive the system. We showed that such analyses, applied to neural activity patterns elicited by viewing various objects, can identify real but subtle and complex features of those objects which are encoded in the brain.

---

Comparisons of object representations in human cortex can give meaningful insight into the neural mechanisms which encode them, and *representational similarity analysis* (RSA) is a popular framework for organizing and analyzing many such comparisons. RSA estimates the *representational geometry* of a (neural) computational system as a matrix of representational similarities (RSM) or dissimilarities (RDM) (1) Two such matrices can then be compared with a *second order isomorphism* to quantify similarity or differences between the two systems – Spearman correlation is commonly used in neuroimaging studies (2, 3). Correlation between RDMs can identify (i) brain regions which similarly represent stimuli, (ii) commonalities in neural codes between species, (iii) computational models which faithfully represent the function of a brain region, and more.

The powerful and flexible machinery of RSA has yielded many successes in neurosciences and neuroimaging in particular – the introductory paper (3) has been cited over 3300 times so far. More pertinent to the current discussion are examples of RSA applied to fMRI vision studies – one study (4) showed that there may be a spectrum of representations of animals in human visual cortex, from most animate to least; another study (5) showed that visual areas represent shape and category to different extents and with interactions; and in (6) it was shown that primates may have a similar neural code for object representations in the IT cortex.

However, there have been several major criticisms of RSA, the strongest being that two computational systems with highly similar RDMs may be carrying out their computations in fundamentally different ways (7, 8). With the primary goal of RSA being to infer whether two computational processes are similar or not, this issue alone may render our inferences from RSA experiments as limited. For example, in Figure 1 we consider a representational space of a torus, i.e. a hollow doughnut. Projected into two dimensions (for instance using multidimensional scaling) the torus becomes a 2D annulus (i.e. a circle with added noise) and representations sampled from the torus project to representations in the annulus. Yet the RDMs of the two sets of representations, one set on the surface of the torus (in 3D) and the other along the 2D annulus are erroneously equated by RSA. This example demonstrates that the central assumption of RSA – that linear distances of matrices sufficiently captures similarity of representational geometry for comparison – is not necessarily true.

**Fig. 1.**
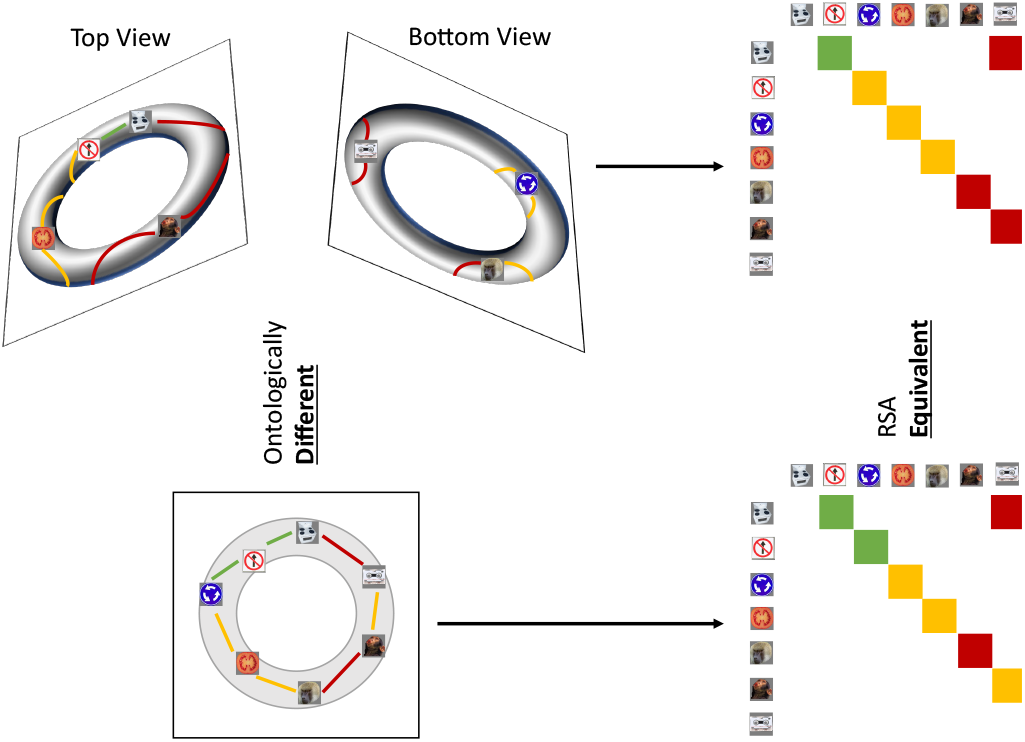
Sampled representations from a torus, projected onto an annulus, and the resulting two RDMs. The stimuli images were obtained from the supplemental information in (9) but were originally introduced in (10), as is the case in later figures. Top left are the top and bottom views of seven sampled representations from the surface of the torus, with colored lines indicating representational distances between adjacent points (green for small distances, yellow for medium and red for large). Bottom left is the projection of these torus representations onto an annulus, with updated representational distances. These distances for both shapes are color-coded in their respective RDMs, which would be considered equivalent by RSA, despite the representational spaces having completely different shapes.

While our example may seem artificial, orientation-selective neurons in V1 have the representational space of a loop (11) and rat grid cells have the representational space of a torus (12). Thus we must use tools that are sensitive to the shape of the representational space of neurophysiological data, or we may err in drawing similarities between two cells (e.g. grid cells and V1 simple cells) based on a simplified model of their representation.

But does this example demonstrate a problem using RDMs to capture representational geometry or rather a problem using correlation as a second-order isomorphism? It has previously been suggested that using non-linear second-order isomorphisms would better account for non-linear geometries (13), and some studies proposed such isomorphisms for analyzing correlation matrices of neurological data (14, 15). A technique called distance correlation (16) has also been shown to be a useful measure of independence in RSA model comparisons (17), being able to capture non-linear dependencies as well as linear ones. These approaches accounted for the nonindependence of pairs of correlation/distance matrix entries, but in the example offered above comparing a torus and its projection onto an annulus, the difference comes from global topology, which implicates distinct causal mechanisms – the annulus contains one periodic phenomenon (captured in one loop) whereas the torus contains two – the major loop and the minor loop which bisects the tube. Such structural features are only detectable when taking into account all dissimilarities together, not just the non-independence of pairs. For example, a Gaussian-distributed cluster, and the same cluster punctured with a small hole in its center, will have similar covariance matrices despite the former being a cluster and the latter being an annulus.

Unfortunately it is not possible in RSA to segment RDMs into features (i.e., sub-components) of their representational geometries. Multidimensional scaling (MDS) (18) has been used to project RDMs into low dimensions for visualization of representational spaces (3), but the projection dimensions are not directly interpretable and are always linear. For example, in (6) RSA found evidence of a shared neural code in primate IT cortex, but MDS embeddings only revealed a blob-like distribution of representations coarsely separated by object category. If segmentation of representational spaces were possible, we could have linked representational similarity to features of the representational spaces (see below), but this is not possible with current RSA methods. In summary, RDMs, and RSA by proxy, do not address complex (i.e, global and not linear) representational geometries and the question of appropriate second-order isomorphism may only be solved once the representational geometry is appropriately captured. The mathematical discipline concerned with studying distance/adjacency between objects in an abstract space is *topology* (19), and tools from computational topology can be applied to multivariate data to derive topological metrics. Structure or shape of data at a local scale can be integrated into global shape descriptors using topological tools, and shape features can then be quantified and analyzed to capture the richness of data structure. Topology has a number of desirable qualities for analyzing representational geometries. For example, the topology of an object does not change when the object is rotated, stretched or reflected (19). Robustness to these transformations would also be expected of representational geometries – the features of neural codes do not depend on the order of the labels of functional units, or the scale of neural activity (20).

The field of *topological data analysis* (TDA) provides practical tools for analyzing data using topology, and TDA has been applied in myriad fields (21). The most established tool in TDA is *persistent homology*, PH (22, 23), which seeks to identify topological features present in data, classified by their dimension. Persistent homology takes as input the distance matrix of a dataset (like an RDM), and identifies structures inherent in the data, such as clusters, loops and voids (which are elements of the H0, H1 and H2 groups respectively). Persistent homology also provides information about the the sizes and density of points belonging to these structures, which can be used to determine which features are significant.

The workflow of the PH algorithm can be seen in Figure 2. The process begins with a sweep through values of a linkage radius – a parameter that defines the extent of the neighbourhood within which two data points would be joined (linked) to form a structure (called the *Vietoris-Rips complex*), and the topological features of these structures at each linkage value are classified as belonging to either H0 (clusters), H1 (loops), H2 (voids), etc. As we sweep through linkage values, features will appear, persist across some range of linkage values, and then disappear – as shown in Figure 2, the points on the loop form that loop only within a certain range of radii *B* and all points eventually fully connect at radius *C*, destroying the loop structure. The linkage values where a feature comes into existence and ceases to exist are called the birth and death values, respectively. For further mathematical details on this process see (22, 23). The birth and death values, along with the feature dimensions, are plotted in persistence diagrams as shown in Figure 3, and points which having death value much larger than their birth value (i.e. their point in the persistence diagram is high above the diagonal line where birth and death are equal) are called “persistent”. A thresholding procedure (24) can then be used to distinguish between persistent, i.e. significant, topological features and non-persistent, i.e. noise, topological features, and an example of this can also be seen in Figure 3. Another useful piece of information that can be extracted for each topological feature is the *representative cycle*, which is a subset of data points in a given topological object. For additional details, see (25).

**Fig. 2.**
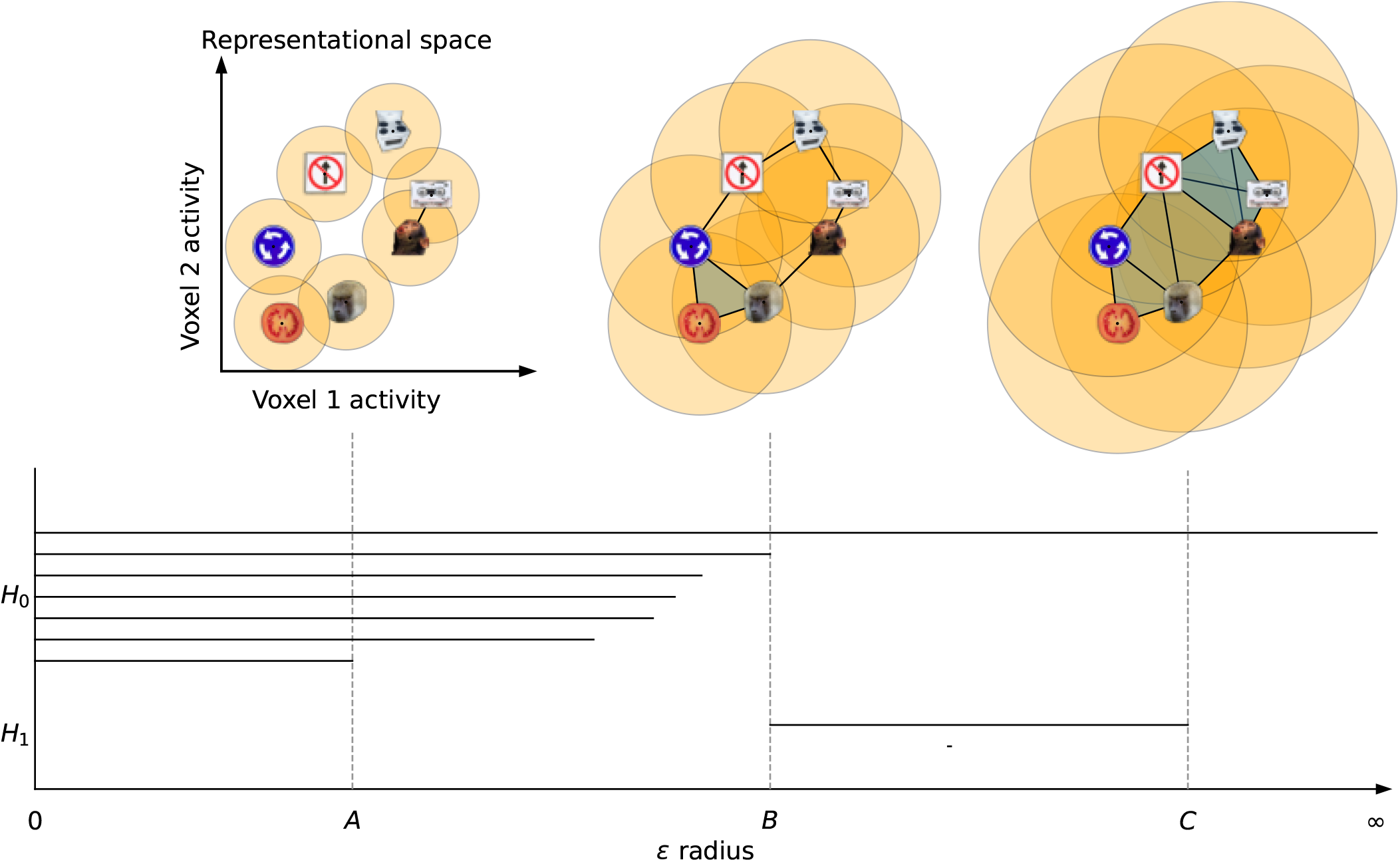
Persistent homology workflow. A linkage radius *ϵ* is increased from 0 and representations (i.e. data points) are connected when their distance is at most *ϵ*, forming Vietoris-Rips complexes. Seven clusters and two loops are present in the dataset, and are tracked by the PH algorithm with each having its own line segment. At linkage radius *A* there are six clusters (since the cassette tape and monkey face are connected, and hence one cluster has died off), while at radius *B* the loop is fully connected (and all components merge into one) and at *C* the loop is filled in (i.e. is no longer a loop).

**Fig. 3.**
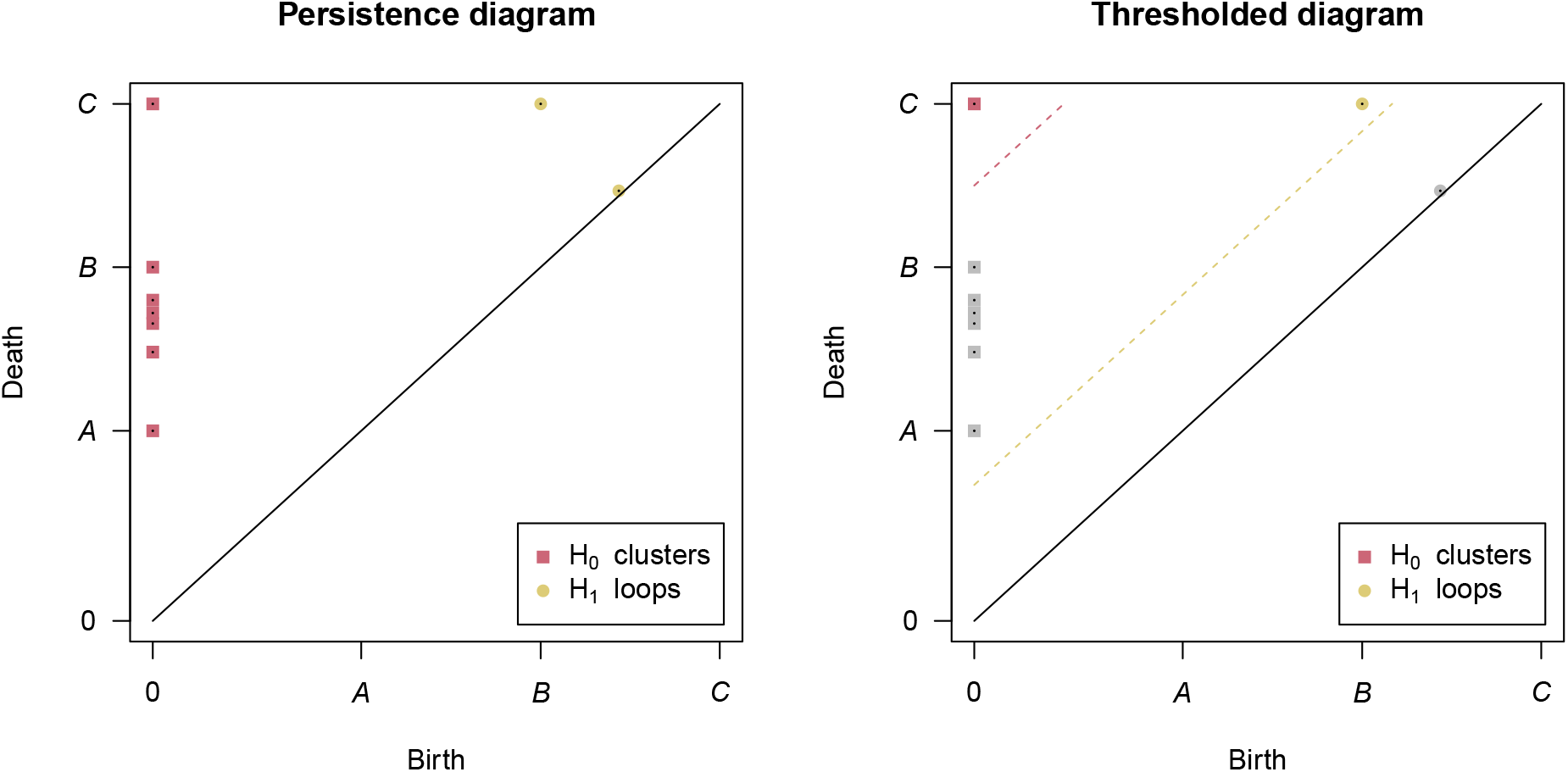
The output persistence diagram of PH run on the example dataset in Figure 2 (left) and an example thresholded diagram (right). In the persistence diagram there are points for each of the seven clusters and two loops one loop is very close to the diagonal line where birth and death are the same, indicating that this loop was very “short-lived”. In the thresholded diagram only one cluster and one loop were significant, indicated by their color and placement above their respective threshold lines.

As the number of sampled points in an object grows, the topological features of the points, recovered by persistent homology, converge to the underlying features (26) even in the presence of noise in the dataset (22). In this sense, increased sampling provides greater validity of the topology of the data space.

It has been demonstrated that persistent homology can detect event-related periodic spatial signals (i.e. spatial loops) in simulated event-related fMRI data (27). A related technique for calculating persistence diagrams called “persistent cohomology” can detect representational space topologies of neural population responses of simulated rat neurons (28). Persistent homology has also been used to find meaningful structure in correlation matrices of spike trains in rat place cells using vectorized summaries of persistence diagrams called Betti curves (29). PH also correctly characterized a low-dimensional neural manifold of mouse behavior analyzing binned spike counts of thalamic neurons (30). The results of these studies suggest that persistence diagrams are a useful tool for characterizing representational spaces, but four properties of diagrams make them particularly well-suited for this task:

- The topological features in persistence diagrams can be identified in their input datasets, thereby allowing us to segment datasets into representational features.
- Persistence diagrams remain consistent under different orderings of the same variables.
- Two persistence diagrams can be meaningfully compared even if their input datasets contained different numbers of data points or variables.
- Persistence diagrams converge as the number of data points in their input datasets grow (26).

Property 1 means that we can uncover topological features of representational geometries, allowing for constraints on the mechanisms implicated while making comparisons between systems more interpretable. For example, a torus and a loop have different numbers of significant loops (two and one respectively), and therefore we could distinguish between RDMs sampled from them. Also, two systems with similar linear aspects in their geometries may perform different calculations and this could be uncovered by investigating their topological sub-structures.

On the other hand, Properties 2 and 3 suggest that using persistent homology to analyze representational geometries may allow for the pooling of data from different studies, even studies with different (but relatable) sets of conditions/stimuli so long as their pooling is defensible and interpretable to the researcher – for examples, studies investigating face processing may use different face conditions, different non-face stimuli, etc, and Properties 2 and 3 of PH allow us to pool results from these studies for stronger inference of representations.

As multiple RDMs can be compared using RSA, we would need an equivalent topological tool to compare multiple persistence diagrams. For second-order isomorphisms of persistence diagrams there exist two main approaches in the literature – for differences, we can use distance calculations (31) and for similarities we can use kernel calculations (32). Since topological features can be comprised of any number of data points (representations), we can capture differences between any number of data points between two representational geometries using these topological second-order isomorphisms. While in regular RSA differences and similarities are essentially opposites (like in the case of correlation and correlation distance), due to the complex nature of persistence diagrams (33) we need specialized and distinct tools to calculate their differences and similarities.

Two typical analyses of RDMs include

- Inference – deciding if two RDMs or two groups of RDMs are similar/different (an important example of which is model comparison), and
- Visualization – using MDS to project an RDM into low dimensions (3).

Similar analyses can be performed with persistence diagrams – differences among sets of persistence diagrams can be found using distance-based permutation approaches as in (34, 35) – and the pairwise distances between multiple persistence diagrams can be used to form an MDS embedding of the diagrams into a low-dimensional space. We have implemented these analytical and inferential tools to carry out TDA on large multivariate datasets (e.g. fMRI) in our software package TDApplied (36). Therefore, the machinery is in place to analyze persistence diagrams computed from RDMs in ways similar to RSA.

We propose a new approach called *representational topology analysis* (RTA) for detecting structures of representational space. In RTA, RDMs are converted to distance matrices (although this is not necessary for correlation dissimilarity matrices; see the methods section) and then analysed with persistent homology, resulting in *persistence diagrams*, i.e. *representational topologies*, that can then be analyzed with topological machine learning and inference methods. Comparing persistence diagrams is preferable to comparing RDMs because the latter do not encode the topology of data space, while the former explicitly represents this information. Representational topology analysis is ideal in conjunction with regular RSA (for inference on linear aspects of data space) in order to make powerful inferences about representational geometry and, by extension, fundamental mechanisms that gave rise to them. Interpretations from RTA are also complementary to interpretations from RSA because in the topological case we can make conclusions about when two representational geometries are different or similar topologically, compared to the regular RSA case where we can only say if two geometries are linearly different or similar. Below, we applied RTA on two datasets and were able to answer questions that regular RSA could not, demonstrating the potential value of representational topology.

## Results

In order to compare RTA with RSA we carried out two studies – the first used data from one of the seminal studies of RSA, (6), and the second used data from a study of shared representations of naturalistic movie viewing across subjects (37, 38).

### Human and Monkey IT Cortex Data

One of the earliest applications of RSA to visual fMRI studies was (6), in which RSA was used to show a common representational code in the primate inferior temporal cortex by comparing fMRI data in humans and electrophysiology data in monkeys (which was collected in the study (10)). But what topological representational features exist in this shared space? Regular RSA cannot segment RDMs to find features of a representational space, and therefore cannot address this question. One of the authors of (6) provided us with the mean RDMs from the group of four humans and the group of two monkeys for the 92 visual stimuli displayed in Figure 4. The stimuli in the experiment were images of various categories, including animals, humans, body parts, naturalistic scenes and objects, and these images can be found in the supplementary data of (6).

**Fig. 4.**
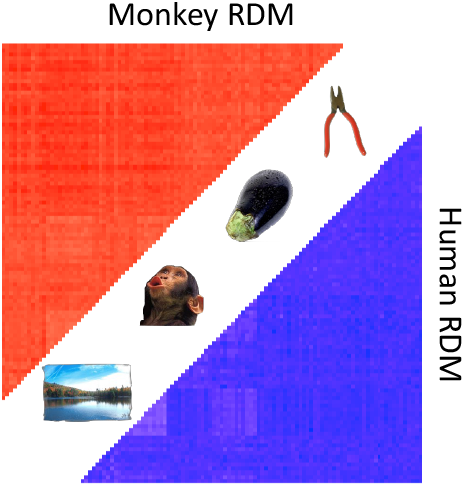
The mean human (bottom right) and monkey (top left) RDMs (each converted ^3^ to a distance matrix using the transformation 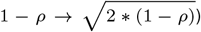. Deeper colors indicate greater representational distances.

A common analysis of persistence diagrams includes identifying the most persistent feature, i.e. the topological feature with greatest difference between their death and birth values – it “lives” longest compared to all other features. We identified the most persistent loop from each of the two diagrams, which we will refer to as the “human loop” and “monkey loop”. For each loop we calculated a representative cycle, i.e. a subset of the 92 stimuli on each loop. Representative cycles are a useful tool for exploring topological features because those features may occupy distinct regions of the data space. For example, imagine a dataset with a loop and a cluster (not touching each other) – all the data points would be used to calculate the data’s persistence diagram but only a subset of the points would lie on the loop. On the other hand, the points that do not lie on the loop, even if they are not part of any other interesting topological structure, can still have immense value in non-topological analyses (for instance in RSA analyses).

A visualization that can help identify data features defined by topological features is the graph defined by a Rips complex, called a *Vietoris-Rips graph* (39) or VR graph for short. At each linkage radius *ϵ* in persistent homology, a set of edges are defined between the data points based on their distances (all the distances *≤ ϵ*), and this defines a graph at each linkage. We visualized the VR graph of the monkey RDM at the linkage radius of the monkey loop birth, and likewise for the human RDM and loop, in Figure 5 to get a sense of the topological structure of the monkey and human RDMs at those linkage scales.

**Fig. 5.**
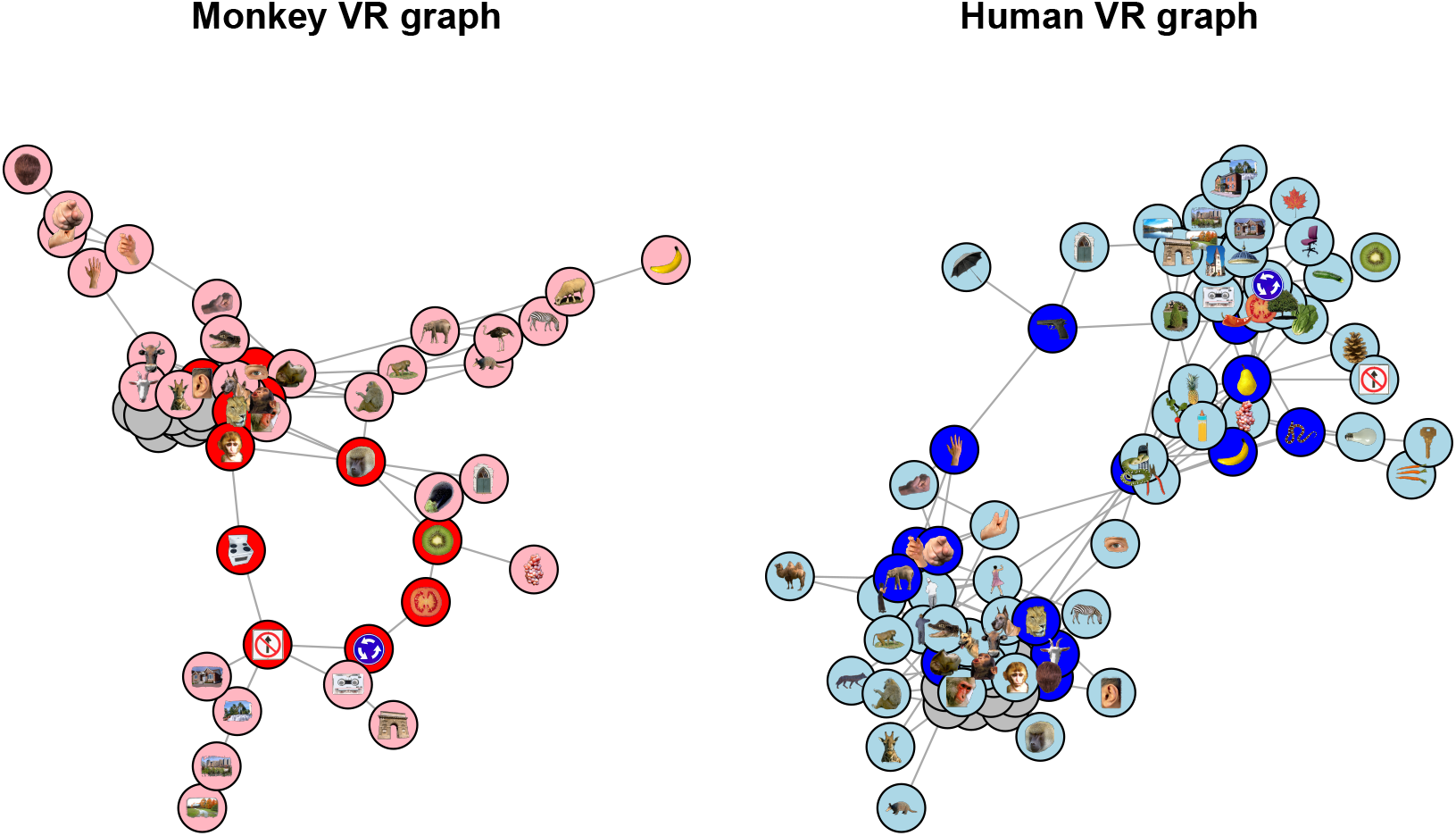
The VR graphs of the monkey RDM (left) and the human RDM (right) at the scales of their respective loop births, with the stimuli in the representative cycles of the two loops highlighted (based on bioRxiv requirements graph nodes which represent face stimuli do not contain their stimuli and are colored gray). The monkey visualization shows a central cluster of animal and monkey faces, from which the loop and two flares (an animal body flair, right, and a hand flair, top left) stem from. From the loop there is also one flair which corresponds to scenery. Only 54 of the 92 stimuli were plotted as these vertices made up the connected component of the VR graph which contained the loop (each of the other 38 stimuli either had no connections to other stimuli or formed small, topologically uninteresting clusters). The human visualization contained 81 of the 92 stimuli, and appears to be two dominant clusters with two paths of sparse connections forming the loop. The clusters are animate objects (left) and inanimate objects (right).

Striking differences occur between the two representational spaces in this view – the monkey VR graph highlights substantially more clustered representations that we can easily label, such as animals, hands, faces, objects, etc., while the human representational spaces appear to be organized into two clusters symmetrically around a loop. That in both cases the representations appear to be lobes organized around a central confluence is intriguing, and may merit greater investigation.

It is worth noting that RSA suggests that the monkey and human representations are highly comparable (6), finding a gross clustering into animate and inanimate objects in both human and monkey spaces, whereas RTA reveals the ways in which they are actually different.

### Naturalistic movie viewing data

In (37), local spatial patterns of BOLD activity in subjects viewing 2D and 3D naturalistic movies (38) were found to be highly conserved across subjects in early visual areas and were modified by region and visual stream – early, ventral and dorsal. It would therefore be expected that group-average topological features differ by region especially for regions in different visual streams. In order to test this hypothesis we analyzed region-level data from (37), constructing timepoint-by-timepoint spatial-pattern correlation distance RDMs (i.e., the correlation distance between time *i* and *j* of the BOLD patterns in each region) and used RTA to characterize the shape of representational space in group average RDMs from certain early, ventral and dorsal regions.

Movie viewing is a naturalistic task that typically induces very similar temporal patterns of activity in a group of subjects (40, 41) and it has recently been shown that this similarity is likely driven by gamma oscillations (42) and is detectable in the spatial patterns in a manner that is more informative of viewing condition (stereoscopic 3D vs mono) than the temporal pattern correlation (43). Here, we used RTA to determine whether the structure of the representational space of spatial patterns over time is different between regions/streams.

To this end we computed group-average region-level significant topological features – loops which survived the thresholding procedure of (24) (see the methods section for details). Significant loops which exist at different scales (i.e. with different birth and death values) would indicate qualitatively different representational structures. We computed the mean RDMs for five regions, across subjects, of higher ventral regions VO1 and VO2, higher dorsal regions PHC1 and PHC2 and the early region V3 in both hemispheres for both 3D movie clips, resulting in 20 RDMs. We chose to analyze only 3D movie data to ensure that there was no confounding effect of stimulus condition in our analysis, and because 3D movies are closer to naturalistic stimuli than 2D movies. We used the bootstrap procedure to identify significant loops, and determined the most persistent loops from the VO regions, PHC regions and V3. The result was three group-average RDMs – one VO RDM, one PHC RDM and one V3 RDM.

In order to compare the three representational spaces, we plotted the VR graphs of the three mean RDMs, at the scale of their respective loop birth values, subsetted to contain only the data points which were in the components of their respective loop representative cycles. Since each graph represents data from one movie across subjects, each graph node represents a TR in its graph’s movie, so each node is plotted with the movie frame five seconds prior to the TR (accounting for the hemodynamic lag). To determine if and where RSA provided a complementary view of these graphs, we projected the three RDMs (subsetted for the TRs in their respective VR graphs) into 2D space using MDS and colored each node in the VR graph by its location in MDS space. We call this novel visualization a *proximity-labelled rips graph* (PLRG for short). Finally, in order to link the PLRG’s back to the raw data we also plotted the movie frame associated with each graph node at the node’s location (in the graph space, not in the MDS space). The results can be seen in Figure 6.

**Fig. 6.**
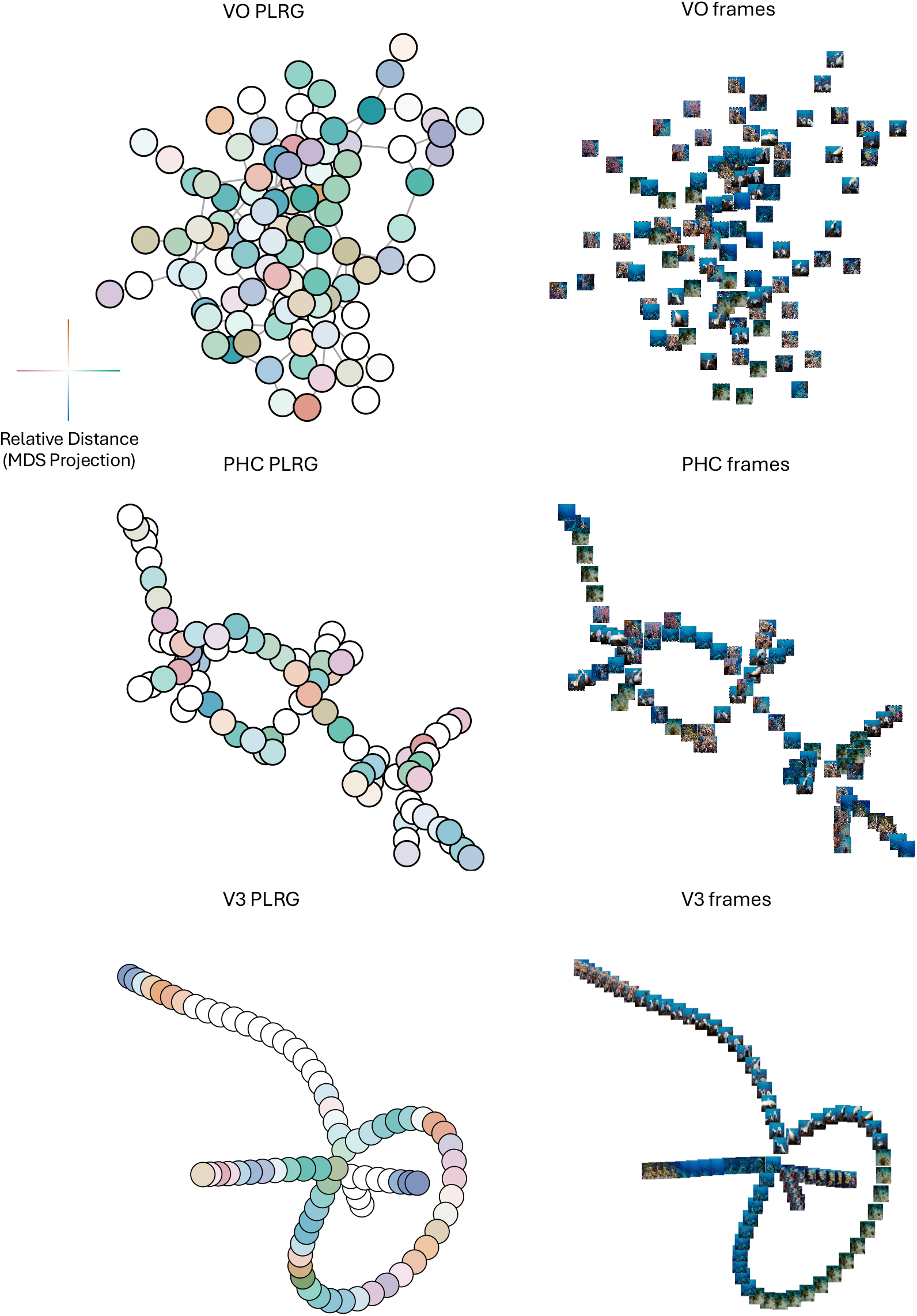
Topologies of mean representational spaces in VO (top row), PHC (middle row) and V3 (bottom row) areas. Left column is the PLRG laid out using a graph-layout algorithm, right column are the frames corresponding to each graph node, plotted at its node’s 2D coordinate in the graph. The color-coding scheme for PLRG nodes, based on MDS coordinates, is displayed to the left of the VO PLRG – the x coordinate determines a horizontal color which is green for positive x-values and purple for negative x-values, and a vertical color which is orange for positive y-values and blue for negative y-values, and two nodes which have similar colors are TR’s with correlated activity patterns, i.e. are nearby in MDS space. PHC and V3 have clearly-defined topologies in their PLRGs, whereas VO has mainly one densely-connected cluster. As well, the lack of color-clustering and smooth color gradients in the VO and PHC PLRG’s indicate that MDS, i.e. RSA, did not capture the graph structure well. V3 on the other hand did exhibit color clustering and gradients, suggesting that there was a stronger relationship between topology and geometry at the loop birth scale. Moreover, the clustering and gradients suggest that some folding of the graph may be appropriate, where nodes which are far apart on the graph with similar colors may actually be proximal in terms of the geometry of data space. The frame visualization of V3 also appeared to most smoothly vary by color and scene type compared to PHC and VO.

The differences in the topological structure between the three regions can be readily appreciated, and these structures were not accounted for by MDS of the RDMs. This illustrates the importance of topological analysis of representational space for inference on similarity.

## Discussion

We demonstrated the potential of topological analysis in identifying representational structures in stimulus-driven fMRI patterns, and showed how this knowledge of the representational geometry can be complementary to standard RSA. Importantly, sensitivity to topological features allows one to find non-linear dimensions in representational spaces, such as the loop we reported for the monkey IT data. This approach goes beyond classic inferential statistics and allows us to have insight into the nature of the mechanisms underlying neural representations.

We first examined two RDMs, one averaged from four human’s IT cortices and one averaged from two monkey’s IT cortices. Unlike in (6) we found that the representational spaces were different because the most persistent loops in the monkey and human representational spaces did not appear to encode the same information – the monkey loop likely encoded a continuous spectrum of change in object category, whereas the human loop was more likely the distal connections between animate and inanimate clusters. Two possible explanations of the differences between the two loops could be that (1) human IT cortex efficiently resolves object category into natural and animate clusters, whereas this distinction is more blurred (i.e. continuous) in monkey IT cortex, or (2) the representational spaces are distinct simply because humans and monkeys can have very different semantic encodings of the same image. The first explanation seems unlikely – the monkey VR graph also had a clear distinction between animate and inanimate objects. The second explanation seems more likely, for example a giraffe and a monkey may have similar representations in humans because they are both animals found in Africa, and in monkeys because they are both non-dangerous creatures – in other words, there is not a one-to-one semantic correspondence. This result is perhaps the best exemplar of the major criticism of RSA described earlier – the two species may be performing very different calculations, and this difference was only detectable using RTA.

We carried out the same analytic approach to a naturalistic movie viewing dataset (37). We analyzed group-mean topological structures in VO, PHC and V3 areas. Our novel proximity-labeled rips graphs of the spaces, at the scale of their most persistent (significant) loop’s birth, were visually very different between the three regions and could not be accounted for by geometry alone – the VO and PHC PLRGs did not have similar colorings of nearby nodes, and the V3 PLRG had clusters of similarly-colored nodes which existed far apart in the graph. By plotting the frames corresponding to 5 seconds prior to each TR over each TR’s graph node, we can see that V3 is likely representing low-level movie features – like object position/movement or scene color, as demonstrated by the many neighboring frames which seem to only differ in the position of objects in the frame or the scene color. On the other hand, there is not a clear division of scene (object) category in the VO PLRG, nor is there a clear relationship between graph structure and object movement/position in the PHC PLRG, but the PHC does exhibit a clear structure (as opposed to VO). In this example RTA was also able to capture aspects of representational spaces which RSA could not.

While RSA provides a “hub” for researchers to integrate data from different modalities, species, etc., it may be limited by the requirement of fixed-size matrices to encode representational geometries and correlations. Because persistent homology

- is invariant under reordering its input data,
- can compare outputs regardless of the number of points it used as input, and
- its output converges as the number of data points grows,

representational topology analysis may be well-suited to compare RDMs across RSA studies which do not have the same stimulus set or set size, building on ideas first proposed in (20). Concrete evidence for this use-case of RTA is hidden in our naturalistic movie analysis – free movie viewing does not follow the traditional task-based experimental design of RSA studies, which is necessary to compute stimulus-stimulus representational dissimilarities, but the flexibility of RTA allowed us to consider each time point as a “stimulus”, the spatial pattern at a time point its “representation”, and carry out principled comparisons of the topological structures which arose across subjects and movies. This means that fMRI datasets (as well as data from other functional neuroimaging modalities) can be compared regardless of their experimental design or duration, which would be particularly interesting for resting-state datasets. Resting state data is characterized by co-fluctuations between distal but functionally-related regions (44–46), which implies the existence of periodic spatio-temporal signals that could be detected with persistent homology – spherical representational topologies have already been identified in restingstate (and naturalistic image viewing) electrophysiological data from V1 in monkeys (11) and this topology could be explained by the interactions between the (periodic) orientation and spatial frequency feature maps. To our knowledge RTA is the first framework that allows comparisons between scans of different duration and study design without temporally collapsing data.

Despite the unique capabilities of RTA, it does have several limitations. Firstly, it is more complicated than regular RSA – there are more computational tools which are needed to carry out a topological analysis. Secondly, RTA is more computationally demanding – persistent homology can be computed quickly with small RDMs (up to around 100 stimuli) in low dimensions, but computing higher-dimensional homology with large RDMs will likely be slower. Similarly, the analysis procedures for persistence diagrams can take time if the persistence diagrams contain many points (although this can be remedied by using the bootstrap procedure to only select significant topological features) or if there are a large number of persistence diagrams (as in a fMRI searchlight analyses). Thirdly, RTA does need a minimum number of stimuli in an experiment to potentially be able to find meaningful topological structure – there are no formal rules, but probably at least ten to find a loop and at least twenty to find a void might be a reasonable assumption. However, in (9) it is suggested that regular RSA performs best when there are many stimuli, so the same would hold for RTA.

Representational topology analysis directly addresses the topology of representational space – an aspect that RSA (as a linear geometric method) cannot. This understanding of representational geometry is useful in that it can reveal nonlinear dimensionality of the representation space which has direct implications for the nature of the input patterns and, by extension, the mechanisms that give rise to those input patterns. In this manner, understanding the topology of representational space provides for novel insights unafforded by existing methods.

## Materials and Methods

### Human vs. Monkey Comparison

We received two RDMs from the authors of (6), one which was the average RDM from four human subject’s 3T fMRI data and the other of which was the average RDM from two monkey subject’s electrode recording data. The entries of the RDMs were (average) correlation distances (i.e. 1 subtract Pearson correlation) between the spatial response patterns of voxels/cells for each pair of stimuli. For more details, see (6). We further transformed the correlation distance values from 1 *− ρ* to 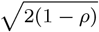 which better satisfy the mathematical notion of distance (36).

We calculated persistent homology of the two RDMs using the R package TDA (47), up to homological dimension 1 (loops), up to the connectivity radius which was the maximum RDM entry, and using the dionysus library functionality (48) to calculate representative cycles (i.e., a subset of the data points that lie on each loop) for the loops. We then computed VR graphs (39) from the two RDMs at the scale of the birth radius of the most persistent loop for each RDM. The stimuli in the two representative cycles were highlighted with deeper colors. The layout of the graph is optimized to project connected nodes nearby each other in 2D space and unconnected nodes further apart, using a graph layout algorithm from the R package igraph (49). We plotted only the graph component which contained the representative cycle nodes. The computation and visualization of the VR graphs was performed by TDApplied.

### Naturalistic movie viewing study

For a detailed account of the data, acquisition and preprocessing of our naturalistic movie viewing analysis, see (37). The study collected 3T fMRI data, with 3mm^3^ voxels, from 55 subjects watching four 5-minute movie clips in one scan (two clips each viewed in both 2D and 3D). The TR was 2 seconds, and the first 1 minute of each movie clip was not analyzed, resulting in 120 TRs of data for each movie clip. Data preprocessing was carried out with the AFNI software (50) and fMRI voxel data was projected onto cortical surface nodes (36002 per hemisphere) with the SUMA (51) and FreeSurfer (52) software packages. Cortical regional boundaries followed the probabilistic atlas from (53).

We chose to solely analyze 3D movie clips in our analysis, in the regions V3, VO and PHC. To calculate an ROI RDM in a hemisphere for a particular movie clip we selected the surface nodes in that hemisphere which were in the ROI (based on atlas boundaries), and computed Pearson correlation between each pair of TRs of the time series activity of all the nodes in that movie. This resulted in a 120×120 representational similarity matrix, which was converted to an RDM by transforming each correlation value *ρ* to the distance value 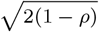. To obtain a group average RDM for each region, movie and hemisphere, we averaged the subject-specific RDMs.

We calculated persistent homology of the RDMs using the R package TDAstats (54), up to homological dimension 1 (loops) and up to the connectivity radius which was the maximum RDM entry. This homology calculation was used in conjunction with the bootstrap procedure (24) in TDApplied to identify significant topological features, and was implemented with 30 bootstrap iterations and significance threshold *α* = 0.1 to avoid over thresholding. The subsetted persistence diagram, according to the bootstrap thresholding procedure, then contained significant group-average region-level topological features (loops). For each region – V3, VO and PHC – we identified the most persistent significant loop out of all its thresholded diagrams, the loop’s birth radius and the RDM it came from. We then used the R package TDA to calculate the representative cycles for those three significant loops from their respective RDMs, by performing the same persistent homology calculation with the dionysus library functionality.

Our novel Proximity-Labelled Rips Graph (PLRG) visualization requires an RDM, the birth scale of a loop and its representative cycle. The nodes of the PLRG graph are the TRs (i.e. spatial patterns) and connections between nodes are determined by the RDM entries which are at most the birth scale (i.e. a PLRG is a VR graph). We plotted only the graph component which contained the representative cycle nodes. Once again the position of the graph nodes in 2D were determined by the igraph package. In order to color the PLRG nodes, the RDM is projected into 2D using the R package stats (55), and the color of each node is determined by the location of its data point in MDS space according to a horizontal color scale (pink (left) to green (right)) and a vertical color scale (blue (bottom) to orange (top)). Outside of calculating the color of each node, the full visualization process of a PLRG is performed by TDApplied.

## ACKNOWLEDGMENTS

We would like to thank Prof. Nikolaus Kriegeskorte for his invaluable feedback on this research. S.B. and R.F. acknowledge funding from the CIHR 2016 grant for cortical mechanisms of 3-D scene and object recognition in the primate brain.

